# Nucleic acid extraction and sequencing from low-biomass synthetic Mars analog soils for in situ life detection

**DOI:** 10.1101/358218

**Authors:** Angel Mojarro, Julie Hachey, Ryan Bailey, Mark Brown, Robert Doebler, Gary Ruvkun, Maria T. Zuber, Christopher E. Carr

## Abstract

Recent studies regarding the origin of life and Mars-Earth meteorite transfer simulations suggest that biological informational polymers, such as nucleic acids (DNA and RNA), have the potential to provide unambiguous evidence of life on Mars. To this end, we are developing a metagenomics-based life-detection instrument which integrates nucleic acid extraction and nanopore sequencing: The Search for Extra-Terrestrial Genomes (SETG). Our goal is to isolate and sequence nucleic acids from extant or preserved life on Mars in order to determine if a particular genetic sequence (1) is distantly-related to life on Earth indicating a shared-ancestry due to lithological exchange, or (2) is unrelated to life on Earth suggesting a convergent origin of life on Mars. In this study, we validate prior work on nucleic acid extraction from cells deposited in Mars analog soils down to microbial concentrations observed in the driest and coldest regions on Earth. In addition, we report low-input nanopore sequencing results equivalent to 1 ppb life-detection sensitivity achieved by employing carrier sequencing, a method of sequencing sub-nanogram DNA in the background of a genomic carrier.

## 1. Introduction

Major strides in understanding the origins of life and meteorite transfer simulations support the notion that life on Mars, if it ever existed, may share a common genesis or perhaps share a common ancestry with life on Earth. Namely, analogous pre-biotic environments (Grotzinger et al., 2014; Johnson et al., 2008; Morris et al., 2010; Ranjan et al., 2017; Ranjan and Sasselov, 2017; Stoker et al., 2010), molecular feedstocks (e.g., hydrogen cyanide) (Adcock et al., 2013; Brack and Pillinger, 1998; Parker et al., 2011), and plausible abiotic reactive pathways predicted on Earth and applicable on Mars may have resulted in parallel origin events in accordance to the RNA-world hypothesis (Benner and Kim, 2015; McKay, 2010; Patel et al., 2015; Powner et al., 2010, 2009; Ritson and Sutherland, 2012; Stairs et al., 2017). This hypothesis suggests that past or present Martian life may have utilized known building blocks (e.g., nucleic acids, sugars, amino acids) and closely resembled life as we know it. Moreover, non-sterilizing lithological exchange between Mars and Earth from impact ejecta produced during the presumed Late-Heavy Bombardment period (Boehnke and Harrison, 2016; Gomes et al., 2005) may have transported viable microbes between planets (Abramov and Mojzsis, 2009; Horneck et al., 2008; Shuster, 2005; Weiss, 2000) resulting in ancestrally related life (Isenbarger et al., 2008).

To test these hypotheses, we are developing the Search for Extra-Terrestrial Genomes (SETG) life-detection instrument for in-situ extraction and nanopore sequencing of nucleic acids (Carr et al., 2017, 2016) from extant or preserved life on Mars. Assuming a convergent adoption of nucleic acids as the unitary solution of genetic information storage and transmission (e.g., DNA or RNA), long-read nanopore sequencing could be capable of detecting non-standard nucleic acids (Carr, 2016; Carr et al., 2017) possibly endemic to Mars (Ranjan et al., 2017). Furthermore, long-read sequencing is of particular significance for taxonomic identification at or below the species level (Benítez-Páez et al., 2016; Brown et al., 2017; Greninger et al., 2015; Quick et al., 2015) which would permit the detection of microbial forward contamination. For instance, in the case of ancestrally related life, comparing sequence data detected on Mars to conserved genes (Harris, 2003; Makarova et al., 1999) on Earth (i.e., the ribosome) (Woese et al., 1975) could unambiguously discriminate contamination from a true life detection (Isenbarger et al., 2008). Conversely, detecting a genetic sequence unlike anything found on Earth (including non-standard bases) could signify a second genesis and perhaps indicate that nucleic acid-based life is likely universal, or at least common.

SETG operates by first extracting and isolating nucleic acids (DNA or RNA) from cells in solid or liquid samples using a modified Claremont BioSolutions (CBIO) solid-phase Purelyse^®^ bacterial genomic DNA extraction kit. Prior studies have utilized OmniLyse, which is Purelyse^®^ without extraction buffers, to lyse cells in an RNA extraction module aboard the International Space Station (Parra et al., 2017). Long-read sequencing is then conducted using the Oxford Nanopore Technologies (ONT) MinION which sequences nucleic acids via ionic current monitoring (Lu et al., 2016) and has been validated to sequence DNA in microgravity (Castro-Wallace et al., 2017; McIntyre et al., 2016), Lunar and Mars gravity (Carr/Zuber *in prep* 2018), and under simulated Mars temperature and pressure (Carr *in prep* 2018).

Our goal for SETG is to be capable of analyzing a variety of environmental samples relevant to the search for life (related or otherwise to life on Earth) on Mars. However, complex soils, especially those containing iron oxides found on Mars (Bell et al., 2000), greatly inhibit nucleic acid extraction from cells due to competitive adsorption onto mineral surfaces (Hurt et al., 2014; Jiang et al., 2012) or destruction due to hydroxyl radicals (Gates, 2009; Imlay and Linn, 1988) during cell lysis (Mojarro et al., 2017b). Additional inhibition may occur in the presence of silicates (Melzak et al., 1996; Trevors, 1996), clays (Greaves and Wilson, 1969), and salts (Henneberger et al., 2006) due to adsorption and or DNA hydrolysis, denaturation and depurination at alkaline to acidic conditions (Gates, 2009). These interactions are further exacerbated in low-biomass environments and by nanopore sequencing, which presently requires substantial (˜ 1 µg) input DNA not likely to be acquired from recalcitrant soils (without amplification) due to limitations related to sequencing efficiency and nanopore longevity (Mojarro et al., 2018). Roughly five in a million nucleobases are sequenced (R9.4 flowcells and chemistry) while nanopores are expended before detection of sub-nanogram input DNA (Mojarro et al., 2017a, 2018) without library preparation modifications.

Extensive literature exists on methods used to mitigate soil-DNA interactions and yield quantifiable DNA from various soils species (e.g., Barton et al., 2006; Direito et al., 2012; Henneberger et al., 2006; Herrera and Cockell, 2007; Lever et al., 2015; Takada-Hoshino and Matsumoto, 2004). Competitive binders or blocking agents such as RNA, random hexamer primers, and skim milk (e.g, Takada-Hoshino and Matsumoto, 2004) are applied to inhibit mineral adsorption sites while dialysis, desalting, or chelation (e.g., Barton et al., 2006) is employed to precipitate or flush soluble metals and salts from soils. Our previous work from Mojarro et al., 2017b focused on adapting the standard Purelyse^®^ bacterial gDNA extraction kit to isolate DNA from Mars analog soils doped with tough-to-lyse spores of *Bacillus subtilis*. In that study, we concluded that with soil-specific mitigation strategies (e.g., desalting and competitive binders), we may be able to achieve adequate DNA extraction yields that are of sufficient purity for downstream sequencing.

In this study, we now focus on validating the modified Purelyse^®^ kit with low-biomass Mars analog soils containing cell densities observed in the driest regions of the Atacama Desert (Navarro-Gonzalez, 2003) and McMurdo Dry Valleys (Goordial et al., 2016). We perform baseline and modified DNA extractions from 50 mg of Mars analog soil containing 10^4^ spores of *B. subtilis* and vegetative *E. coli* cells, respectively. These experiments stress the importance of the mitigation strategies used to reduce detrimental soil-DNA interactions (e.g., adsorption to mineral surfaces, DNA destruction) and conclude in the development of a ‘near-universal’ extraction protocol for low-biomass Mars analog soils. As in our previous experiments (Carr et al., 2017, 2016), we require DNA extraction yields of at least 5% in order to achieve a minimum sensitivity target of 1 ppb for life-detection (i.e., 10^4^ cells in 50 mg of soil).

Lastly, we investigate low-input nanopore sequencing by experimenting with 2 pg of purified *B. subtilis* DNA equivalent to 5% DNA yield from 10^4^ spores. For many environmental samples, the total extractable DNA is far below the current input requirements of nanopore sequencing, preventing sample-to-sequence metagenomics from low-biomass or recalcitrant samples. On Mars, whole genome amplification could result in biasing microbial population results (Sabina and Leamon, 2015) while targeted amplicon sequencing such as 16S rRNA would assume a shared ancestry between Mars-Earth and could reduce taxonomic resolution (Poretsky et al., 2014). The absence of amplification is especially important in the case where non-standard bases have been incorporated into the Martian genome as primers may not function or amplification could theoretically mask an alien signal due to promiscuous base-pairing resulting in an ATGC sequence (Pezo et al., 2014). Here we address these problems by employing carrier sequencing, a method to sequence low-input DNA by preparing the target DNA with a genomic carrier to achieve ideal library preparation and sequencing stoichiometry without amplification (Mojarro et al., 2018; Raley et al., 2014). We then use CarrierSeq (Mojarro et al., 2018) a sequence analysis script used to identify the low-input target reads from the genomic carrier, and analyze the results in the context of life detection.

## 2. Materials and Methods

### 2.1. Mars analog soils

A total of six Mars analog soils and one lunar basalt analog (Orbitec, JSC-1A) (McKay et al., 1993) were utilized to develop our extraction protocols. Five Mars analog soils were produced in accordance to *in situ* mineralogical and geochemical measurements collected by rover and lander missions as described by Schuerger et al., 2017, 2012. These soils represent various potentially habitable paleoenvironments of astrobiological significance, for instance, ancient hydrothermal salt deposits in Gusev crater (Ming et al., 2006; Ruff and Farmer, 2016). The five soils represent (1) the highly-oxidized global aeolian dust (Bell et al., 2000; Ming et al., 2008a), (2) salt-rich (Burroughs subclass) soils of Gusev Crater (Ming et al., 2006; Morris et al., 2008), (3) jarosite-containing acidic soils (Paso Robles class) of Meridiani Planum (Klingelhofer, 2004; Ming et al., 2008a; Morris et al., 2006), (4) carbonate-rich alkaline Viking Lander soils (Clark et al., 1982; Wänke et al., 2001), and (5) perchlorate-rich Phoenix Lander soils (Ming et al., 2008b). In addition, we include a commercially available (6) aeolain spectral analog (Orbitec, JSC-Mars-1A) (Allen et al., 1998). We henceforth refer to these Mars analog soils as aeolian, salt, acid, alkaline, perchlorate, and JSC, respectively. The lunar analog represents our unaltered soil control and is referred to as basalt. Details concerning the exact methods used to synthesize these soils can be found in Schuerger et al., 2012 and in Mojarro et al., 2017b.

### 2.2. Bacillus subtilis spores

Spore suspensions of *Bacillus subtilis* (ATCC 6633) similar to those used in clean-room sterilization effectiveness (Friedline et al., 2015) and bacteriostasis testing under Mars-like conditions (Kerney and Schuerger, 2011; Schuerger et al., 2017, 2003) were acquired from Crosstex (Part# SBS-08) to represent a worst case DNA extraction scenario of a tough-to-lyse organism. Spore suspension were DNAse-treated (New England Biolabs, M0303L) in order to remove any extracellular DNA and counted using a viable spore assay on lysogeny broth agar plates (Mojarro et al., 2017b). We assume a single genome copy per DNAse-treated spore of *B. subtilis* for calculating DNA yield (DNA_out_/DNA_in_).

### 2.3. Vegetative Escherichia coli cells

Vegetative *E. coli* (OP50) cell cultures were grown overnight in flasks containing 200 mL of lysogeny broth medium inside an Innova^®^ 44 incubator at 37° C and 200 rpm. After 12 hours, we measured optical density (OD600) on a DeNovix DS-11+ spectrophotometer, pelleted, and re-suspended culture aliquots in phosphate-buffered solution (Thermo Fisher, 10010023). An estimated 1000 *E. coli* cells then underwent colony droplet digital polymerase chain reaction (ddPCR) with single copy *metG* (Wang and Wood, 2011) primers for absolute genome quantitation. In contrast to spores which contain a single genome copy, copy-number variation may exist in vegetative cells depending on growth stage (Pecoraro et al., 2011; Skarstad et al., 1986). Therefore, ddPCR results allow us to correct for genome copy-number variation (e.g., 2000 genome copies / 1000 cells) and calculate DNA yield similar to *B. subtilis* (DNA_out_/DNA_in_).

### 2.4. DNA yield quantitation

Absolute DNA yield quantitation is possible through ddPCR paired with single copy primers. In contrast to traditional PCR, ddPCR fractionates one reaction into ˜20,000 water-oil emulsion nanodroplets each capable of discrete amplification. When a sample is adequately dilute, Poisson statistics can precisely determine genome copy numbers independent of a standard curve (Brunetto et al., 2014; Yang et al., 2014). Paired with single copy primers and the presence of a DNA-specific fluorescent dye (EvaGreen), the results of a ddPCR reaction are either relatively weakly-fluorescent droplets containing primer dimer or relatively highly-fluorescent droplets containing an amplified product. We then equate one highly-fluorescent droplet to one genome copy. DNA extracted from *B. subtilis* spores was amplified with *spaC* primers (forward: TGA GGA AGG ATG GGA CGA CA, reverse: AAC AGA TTG CTG CCA GTC CA) (Hu et al., 2004) while DNA extracted from vegetative *E. coli cells* was amplified with *metG* primers (forward: GGT GGA AGC CTC TAA AGA AGA AG, reverse: AGC AGT TTG TCA GAA CCT TCA AC) (Wang and Wood, 2011). Two ddPCR reactions per DNA extraction (one per elution) were performed and quantified on a BioRad QX200 ddPCR system. A total of 5 µL from each extraction elution was prepared in a 20 µL final reaction volume containing 3 µL of molecular grade water, 10 µL of QX200 EvaGreen SuperMix (BioRad, 186-4033), 1 µL of 3.3 µM forward primer, and 1 µL of 3.3 µM reverse primer. All ddPCR reactions were prepared by an Andrew Alliance liquid handling robot within an AirClean 600 PCR Workstation while nanodroplets were generated using the automated QX200 AutoDG system. Thermocycling conditions were as follows: (1) 95° C for 5 minutes, (2) 95° C for 30 seconds, (3) 60° C for 1 minute, (4) Repeat steps 2-3 40 times, (5) 4° C for 5 minutes, (6) 90° C for 5 minutes, (7) hold at 4° C until ready to measure.

### 2.5. Purelyse^®^ standard extraction protocol

The Purelyse^®^ bacterial gDNA extraction kit is a miniature (dime-sized) and battery-powered bead-beating lysis device capable of solid-phase nucleic acid extraction without the need for centrifugation. Purelyse^®^ kits work by shearing cells open at 30,000 rpm in the presence of a supplied low-pH (˜3.5 pH) binding buffer which promotes the binding of negatively charged polymers to the surface of specialty oxide ceramic microbeads (˜100 µm). The microbeads are then washed with a lower concentration binding buffer and DNA is eluted in a low-salt and high-pH (˜8.0 pH) elution buffer. The standard extraction protocol for cell cultures dictates: (1) 2-minute lysis with 1x binding buffer solution (100 µL of 8x binding buffer and 700 µL of molecular-grade water), (2) 45-second wash with 3 mL of 0.25x binding buffer wash solution (˜100 µL of 8x binding buffer and 2.9 mL of molecular-grade water), (3) two 1-minute elutions with 200 µL of 1x elution buffer all at 6V (equivalent to 4 AAA batteries).

### 2.6. Baseline extractions of spore and vegetative DNA from water

Two dilution series containing 10^8^, 10^6^, and 10^4^ spores of *B. subtilis* and equivalent vegetative *E. coli* cells in water were processed using the standard extraction protocol. The *E. coli* dilution series originated from a single culture prepared in phosphate-buffered solution and DNA yield was corrected for genome copy-number variation using ddPCR.

### 2.7. Unmodified extractions of spore and vegetative DNA from Mars analog soils

An estimated 1.6 x 10^4^ spores (about 70 pg of DNA) of *B. subtilis* were deposited on 50 mg of Mars analog soil and processed following the standard extraction protocol. For vegetative *E. coli* extractions, we utilized OD600 measurements to deposit approximately 1.6 x 10^4^ cells on 50 mg of Mars analog soil or equivalent to spore concentrations. However, as *E. coli* genome copy number may vary (Pecoraro et al., 2011; Skarstad et al., 1986) each *E. coli* extraction from a Mars analog included accompanying colony ddPCR results used for correcting genome copy-number variation.

### 2.8. Modified extractions of spore and vegetative DNA from Mars analog soils

To parallel the unmodified extractions, an estimated 1.6 x10^4^ spores of *B. subtilis* and vegetative cells of *E. coli* were deposited on 50 mg of Mars analog soil, respectively. All samples were processed using the ‘near-universal’ modified extraction protocol: (1) Suspend sample in 800 µL of 8x binding buffer and vortex gently for 30 seconds, (2) desalt in a single 100K Amincon Ultra column (Z740183), (3) re-suspend the sample in 400 µL of molecular grade water and 400 µL of 8x binding buffer, (4) add 4-6 µg of random hexamer primers (Promega, C1181) and vortex gently for 30 seconds, (5) lyse cells and bind DNA with Purelyse^®^ at 6.5 V for 2 minutes, (6) wash with 2.5 mL of 1x binding buffer at 1.5 V for 1 minute, (7) elute DNA with 200 µL of heated elution buffer to 70° C for 1 minute, (8) repeat step 7 for second elution. Basalt and alkaline samples contained 4 µg of random hexamer primers while JSC, acid, salt, aeolian, and perchlorate samples contained 6 µg.

### 2.9. Nanopore sequencing

The Oxford Nanopore Technologies (ONT) MinION sequencer and flowcell perform single-strand DNA sequencing through monitoring changes in an ionic current produced by the translocation of k-mers through a nano-sized pore (Lu et al., 2016). Nanopore sequencing encompasses library preparation, sequencing, and basecalling. First, library preparation is the process by which genomic DNA is converted into a readable format for the sequencer. This includes adding tethers and a motor protein onto double-stranded DNA which guide and regulate the translocation rate (R9.4, 450 bp/s) through a nanonopore. Second, the prepared library is loaded onto the flowcell and sequencing is initiated. Up to 512 sequencing channels (pores) can be monitored and raw current (translocation) events are recorded. Lastly, the raw events are then basecalled into corresponding nucleobase assignments or oligomer sequences.

### 2.10. Low-input sequencing library preparation

The sequencing library was prepared following a modified one-direction (1D) Lambda control experiment protocol (Oxford Nanopore Technologies, SQK-LSK108). The ligation-based sequencing kit advises shearing 1 µg of Lambda genomic DNA to 8 kb fragments and spiking the sheared gDNA with a 3.6 kb positive control (Lambda genome 3’-end amplicon) in order to distinguish library preparation from sequencing failures. However, after validating our library preparation proficiency, we substituted 2 pg of *B. subtilis* DNA purified with Purelyse^®^ in lieu of the 3.6 kb positive control and replicated the Lambda control protocol without additional modifications. This substitution results in a low-input carrier library with ideal stoichiometry (Mojarro et al., 2018).

### 2.11. Sequencing and basecalling

The low-input carrier library was sequenced using a MinION Mark-1B sequencer and R9.4 spot-on flowcell for 48 hours on an Apple iMac desktop computer running MinKNOW 1.5.18. The resulting raw nanopore reads were basecalled using ONT’s Albacore 1.10 offline basecaller.

### 2.12. Sequence analysis

All reads located in the Albacore workstation folder were compiled into a single fastq file and mapped directly to the *B. subtilis* reference genome using bwa (Li, 2013). In the context of an unknown sample, all reads were then processed with CarrierSeq (Mojarro et al., 2018), a sequence analysis script for carrier sequencing. CarrierSeq works by identifying all reads not belonging to the carrier (Lambda), applies quality-control filters, and implements a Poisson test to identify likely nanopore sequencing artifacts known as high-quality noise reads (HQNRs) which presumably originate from malfunctioning nanopores. The final subset or ‘target reads’ should therefore only contain *B. subtilis* and likely contamination (Mojarro et al., 2018).

## 3. Results

### 3.1. Baseline extractions of spore and vegetative cell DNA from water

Water extractions of spore DNA remained consistent throughout the dilution series. At 10^8^, 10^6^, and 10^4^ spores, yields were 14.7%, 9.7%, and 13.3% (Figure 1). In contrast, extractions of vegetative cell DNA decreased with cell concentrations. At 10^8^, 10^6^, and 10^4^ vegetative cells, yields were 40%, 21.8%, and 13.2% (Figure 1).

**Figure 1:**
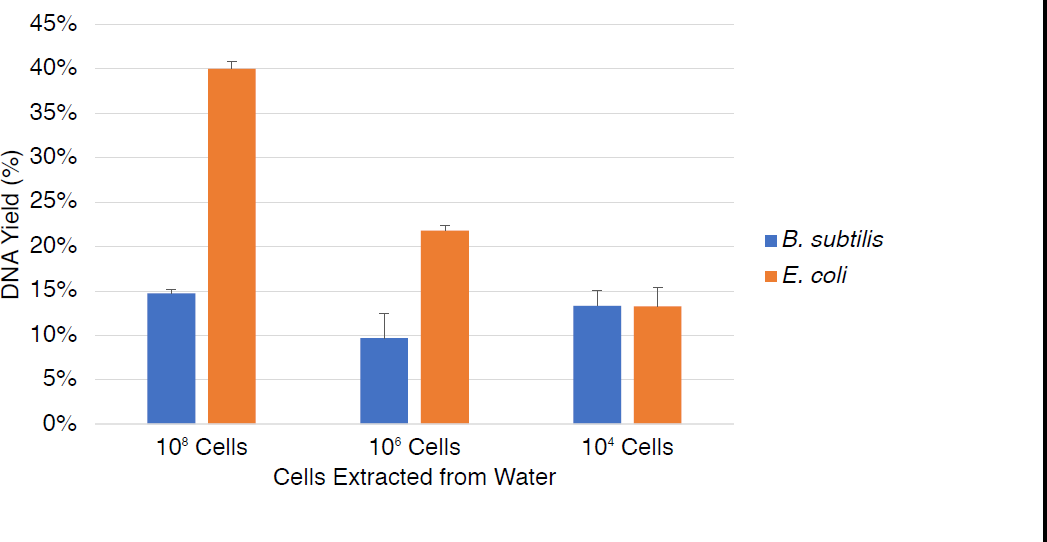
Baseline extractions of spore and vegetative cell DNA from water. Two dilution series containing 10^8^, 10^6^, and 10^4^ spores of *B. subtilis* and equivalent vegetative *E. coli cells* were processed using the standard extraction protocol. Water extractions of spore DNA remained consistent throughout the dilution series. In contrast, extractions of vegetative cell DNA decreased with cell concentrations (standard error shown, *n* = 3).

### 3.2. Unmodified extractions of spore and vegetative cell DNA from Mars analog soils

No detectable DNA was measured from any Mars analog soil extraction containing either spores or vegetative cells with the exception of the basalt and alkaline soils. Unmodified spore DNA extractions yielded 7.1% for basalt and 8% for alkaline while vegetative DNA extractions yielded 4.7% for basalt and 5.5% for alkaline (Figure 2).

**Figure 2:**
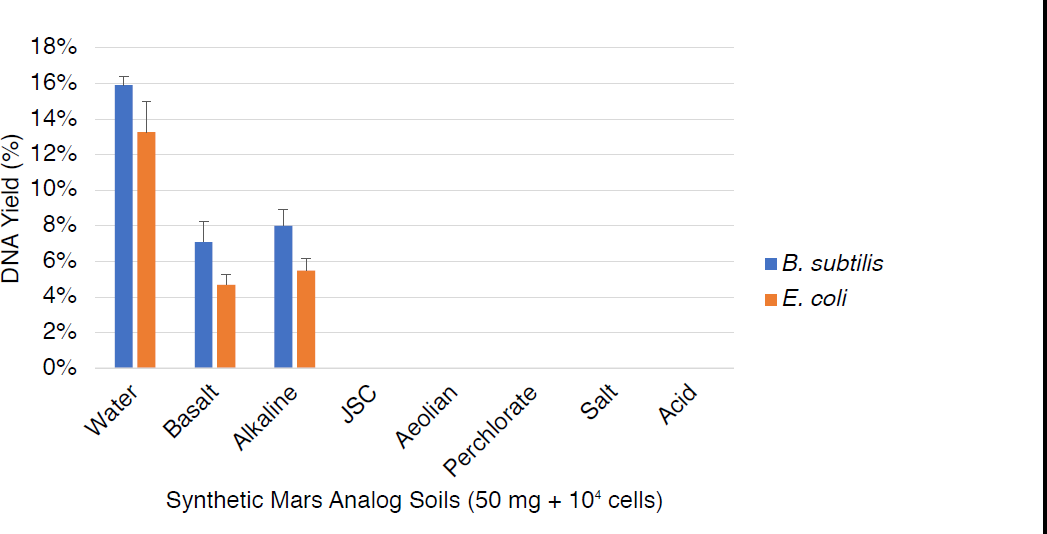
Unmodified extractions of spore and vegetative cell DNA from Mars analog soils. An estimated 1.6 x 10^4^ spores (about 70 pg of DNA) of *B. subtilis* and equivalent vegetative cells of *E. coli* were deposited on 50 mg of Mars analog soil and processed following the standard extraction protocol. No detectable DNA was measured from any Mars analog soil extraction containing either spores or vegetative cells with the exception of the basalt and alkaline soils (standard error shown, *n* = 3).

### 3.3. Modified extractions of spore and vegetative cell DNA from Mars analog soils

All modified Mars analog soil extractions of spore DNA achieved our 5% requirement using the ‘near-universal’ protocol. DNA yields were modestly increased for alkaline and basalt samples while aeolian, salt, acid, perchlorate, and JSC samples increased significantly from undetectable amounts. The spore results were: basalt 9.2%, alkaline 10.7%, aeolian 6.1%, salt 6.7%, acid 5.1%, perchlorate 7.3%, JSC 7.8% (Figure 3). In contrast, several vegetative cell DNA extractions, e.g., JSC, aeolian, and perchlorate, failed to achieve 5% DNA yield. Similar to the spore extractions, yields for alkaline and basalt samples received a moderate increase while the remaining samples increased from undetectable amounts. The vegetative cell results were: basalt 13.1%, alkaline 15.3%, aeolian 4.4%, salt 13.3%, acid 8.0%, perchlorate 4.5%, JSC 2.0% (Figure 3).

**Figure 3:**
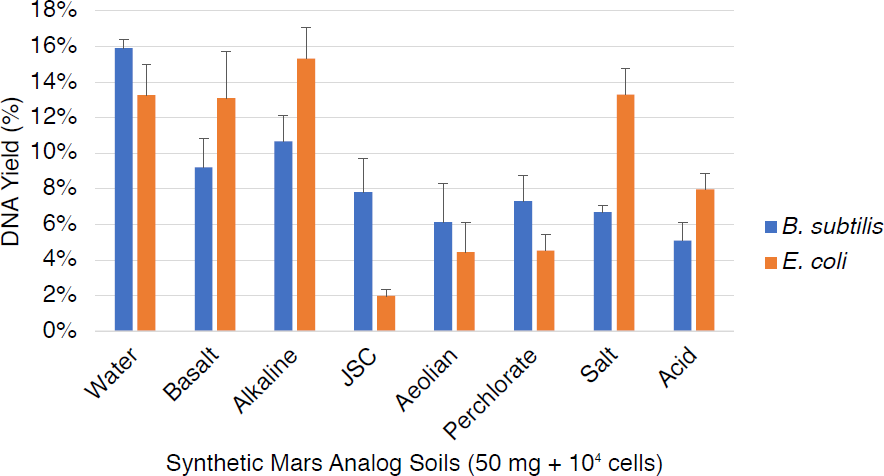
Modified extractions of spore and vegetative DNA from Mars analog soils. All DNA yields from 10^4^ vegetative cells and spores in 50 mg of Mars analog soils were increased with the ‘near-universal’ extraction protocol. Our results indicate that a combination of desalting and completive binders is an adequate strategy for achieving our extraction goals (standard error shown, *n* = 3).

### 3.4. Nanopore sequencing

From the resulting 48-hours of sequencing, we detected a total of 8.7 gigabases or 1.3 million reads. Exactly 1,260,661 reads mapped to Lambda while 5 reads mapped to *B. subtilis*. Given the sequencing libraries Lambda to *B. subtilis* mass ratio, we expected approximately 2 *B. subtilis* reads per 1,000,000 Lambda reads. However, we detected ˜4 *B. subtilis* reads per 1,000,000 Lambda reads. Assuming Lambda is the only known sequence, the CarrierSeq analysis (Q = 9, p = 0.05, default values) isolated 29 ‘target reads’ (Table 1) that were identified using the NCBI blastn algorithm: 5 *B. subtilis* reads, 13 *Homo sapiens* reads, 1 *Klebsiella pneumoniae* read, 1 Human mastadenovirus read, 1 Human DNA sequence (not able to be resolved to *Homo sapiens*), 1 *Streptococcus sp.* read, 1 *Staphylococcus epidermis* reads, and 6 unidentified high-quality noise reads (HQNRs) (Table 2). The basecalled and unanalyzed sequencing fastq files can be downloaded from FigShare: https://doi.org/10.6084/m9.figshare.5471824.v2 or from the NCBI SRA database: [fast5.tar.gz upload pending and available upon request]

**Table 1.**
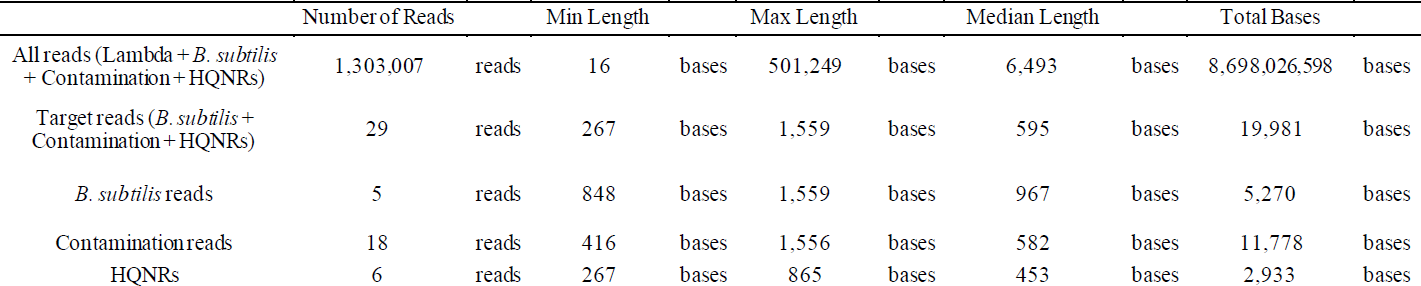
Low-input carrier sequencing metrics.

**Table 2:**
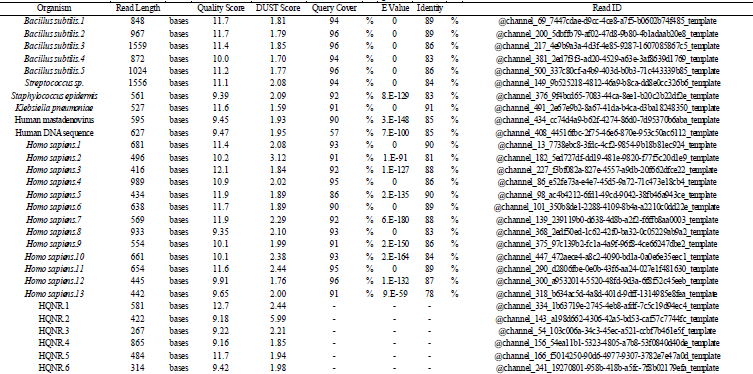
Target Reads.

## 4. Discussion

### 4.1. Extraction of DNA from synthetic Mars analog soils

The extractions from spore DNA in water and soils at 10^4^ spores yielded similar results to our prior experiments at 10^8^ spores used to develop the initial modified protocol (Mojarro et al., 2017b). We previously hypothesized that low DNA yields from *B. subtilis* was generally due to small acid-soluble proteins (SASPs) which bind to the DNA phosphate backbone and furnish protection from heat, salts, desiccation, and UV radiation in the spore state (Moeller et al., 2012, 2009). In short, SASPs could inhibit interactions between the phosphate backbone and the Purelyse^®^ oxide ceramic microbeads, which would ideally attract negatively charged polymers (i.e., DNA). In Mojarro et al., 2017b we seemingly validated this hypothesis by increasing spore DNA yields in water from 15% to 43% using a proof of concept protein separation binding buffer/phenol cocktail. We therefore intuitively expected the DNA extraction yields from vegetative *E. coli* cells, which presumably do not contain SASPs, to be higher than *B. subtilis* spores. Extractions of spore DNA from our water control consistently yielded ˜15% at all cell concentrations while extractions of vegetative DNA began with 40% at 10^8^ cells and decreased to 13.3% at 10^4^ (Figure 1). These extractions share the exact parameters across cell lysis and DNA elution voltage, time, buffer volumes, and approximate cell concentrations with the exception of cells being either spores or vegetative. We are presently unable to offer a definitive explanation for the decrease in vegetative DNA with decreasing cell concentration and the apparent convergence of DNA yields at 10^4^ cells (Figure 1). Perhaps (1) cell concentration may affect DNA yield, (2) binding of nucleic acids to the Purelyse^®^ oxide ceramic microbeads is proportional to the concentration of lysed DNA, (3) vegetative DNA is more prone to destruction (e.g., hydrolysis, etc) in solution without SASPs, or (4) vegetative DNA has a stronger affinity towards the Purelyse^®^ oxide ceramic microbeads and/or soil. Further work is ongoing in order to investigate this inconsistency between spore and vegetative DNA yields.

In regard to the Mars analog soil extractions, prior work from Mojarro et al., 2017b had suggested that both alkaline and basalt soils would exert mild soil-DNA interactions due to adsorption effects by silicates (Melzak et al., 1996; Trevors, 1996; Zhou et al., 1996) while highly-oxidized iron sulfate-containing Mars analog soils would undoubtedly destroy most DNA in an unmodified extraction (Figure 2). Nevertheless, once disruptive metals/cations (e.g., Fe^3+,2+^, Ca^2+^) were flushed and mineral adsorption sites were coated with competitive binders, DNA yields of at least 5% were achieved in all soils containing spores of *B. subtilis* (Figure 3). Extractions from vegetative *E. coli* cells were only capable of achieving a 5% yield within their standard errors with the exception of JSC (Figure 3). Comparing the mean spore and vegetative DNA yields for JSC, aeolian, and perchlorate soils suggests that perhaps DNA bound by SASPs increases resistance to destruction post-cell lysis and pre-binding to the Purelyse^®^ oxide ceramic beads. In such a case, it appears that DNA is more susceptible to damage from free radicals in Mars soils containing Fe^3+,2+^ species (JSC, aeolian, perchlorate) than from hydrolysis, depurination and/or denaturation in acidic to alkaline soils (alkaline, salt, acid) (Figure 3).

### 4.2 Ancient DNA on Mars

The relatively stable cryosphere on Mars since the Late-Noachian period (Head and Marchant, 2014) makes it the ideal location for ancient DNA preservation. On Earth, the estimated half-life of DNA at −25° C is roughly on the order of 10^7^ years (Millar and Lambert, 2013). However, local and global natural climate variability due to orbital cycles (Imbrie et al., 1992) and tectonics (Raymo and Ruddiman, 1992) limit the extent of cold environments favorable towards ancient DNA preservation on geologic timescales. Today, the average temperature at Gale Crater is −48 C (Haberle et al., 2014) while similar conditions may have persisted since the Amazonian period (Head and Marchant, 2014) theoretically permitting the preservation on DNA beyond 10 million years. However, the likelihood of ancient DNA on the surface of Mars greatly decreases once we consider the effect of UV, cosmic ray exposure, and radioactive decay on nucleic acids (Hassler et al., 2014; Kminek et al., 2003; Kminek and Bada, 2006). Ancient DNA is prone to damage (e.g., hydrolysis, depurination) and fragmentation (Dabney et al., 2013; Millar and Lambert, 2013) once it is no longer actively repaired in a biologic system. In the unlikely case that nucleic acids are preserved and accessible at the surface, we then risk the possibility of destroying them once soils are hydrated during an extraction if they are not protected within cells (Gates, 2009; Mojarro et al., 2017b). The primary target for SETG is therefore extant or recently dead cells encapsulating nucleic acids that may survive the sample prep processes and yield long strands useful for taxonomic identification.

### 4.3. Low-input Nanopore sequencing and contamination detection

A great advantage of nanopore sequencing is the ability to produce long reads capable of taxonomic identification (Brown et al., 2017; Goordial et al., 2017; Greninger et al., 2015; Quick et al., 2015). This has allowed us to directly map all basecalled reads to the reference genome and identify the 5 belonging to *B. subtilis* without further processing. However, in a true life detection or unknown scenario, there must be a way to filter carrier (Lambda), low-quality, and low-complexity reads in order to identify those belonging to the unknown organisms. As mentioned earlier, we used CarrierSeq which was developed in order to analyze all carrier sequencing reads and identify the ‘unknowns’ in this context. This analysis identified 29 reads which did not map to Lambda, were not considered low-quality (Q < 9, Albacore passing was Q > 6 at the time of analysis, currently Q > 7) or low-complexity (Morgulis et al., 2006), and did not originate from overly active “bad pores” that are known to produce spurious reads which we referred to as high-quality noise reads (HQNRs) in Mojarro et al., 2018. These include the 5 *B. subtilis* reads detected by mapping, 6 HQNRs that originated from “good pores”, and most interestingly, 18 *Homo sapiens* and human microflora contamination reads (Table 2). Although we cannot recollect an exact point of contamination, this work was conducted in an open lab, with non-sterile gloves, and in an open bench without any non-standard precautions (e.g., routine sporicidal/bleaching was conducted between sets of extractions).

The close abundance of contamination to *B. subtilis* reads possibly suggest comparable low levels of contamination (2 pg of *B. subtilis*), although, there is no certain strategy to ascertain the true initial contamination quantities. We instead see this as an opportunity to highlight the effectiveness of nanopore sequencing to discriminate between true detection and contamination. Particularly in the context of planetary protection where carrier sequencing could be employed as an effective method of detecting low-levels of microbial contamination on the surface of spacecrafts (Rummel and Conley, 2018). The presence of HQNRs, however, complicate the certainty between a true unknown read and noise. In Mojarro et al., 2018 we proposed that HNQRs could possibly be the result of pore blockages from macromolecules (e.g., proteins) that may conceivably produce false signals. To date we have not produced a fully satisfactory approach to identify and discard such reads. Identifying HQNRs through k-mer matching (Menzel et al., 2016) and via the NCBI blastn algorithm is presently the best approach, while plotting DUST (Morgulis et al., 2006), quality (Phred) score, and read length and statistical tests not reveal any obvious relationships (Figure 4, 5). Recent work by Pontefract et al., 2018 on the failure modes of nanopore sequencing has suggested that improvements to basecalling algorithms, library preparation, and sequencing chemistries have suppressed the occurrence of these spurious reads. However, as ONT matures its sequencing technology, more robust testing under simulated unknown sample conditions is required in order to confidently identify HQNRs (if they are still an issue) where these artefactual reads may complicate the identification of a new organism, or specifically, extraterrestrial life.

**Figure 4:**
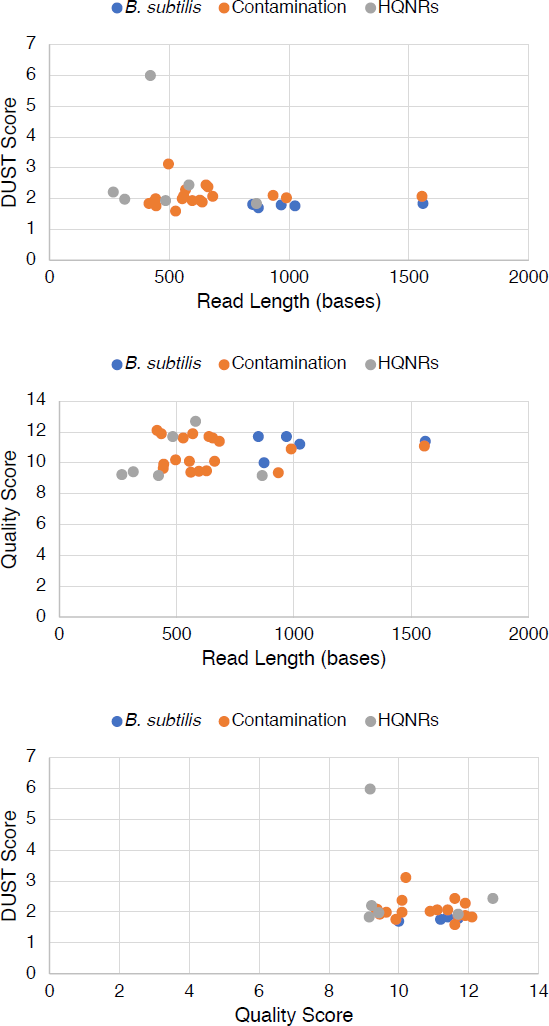
Cross-plots of target reads identified by CarrierSeq displaying read length, quality score, and DUST score. There does not appear to be any separation of HQNRs and *B. subtilis* or contamination reads. We are presently only able to identify HQNRs through k-mer matching or via the NCBI blastn algorithm.

**Figure 5:**
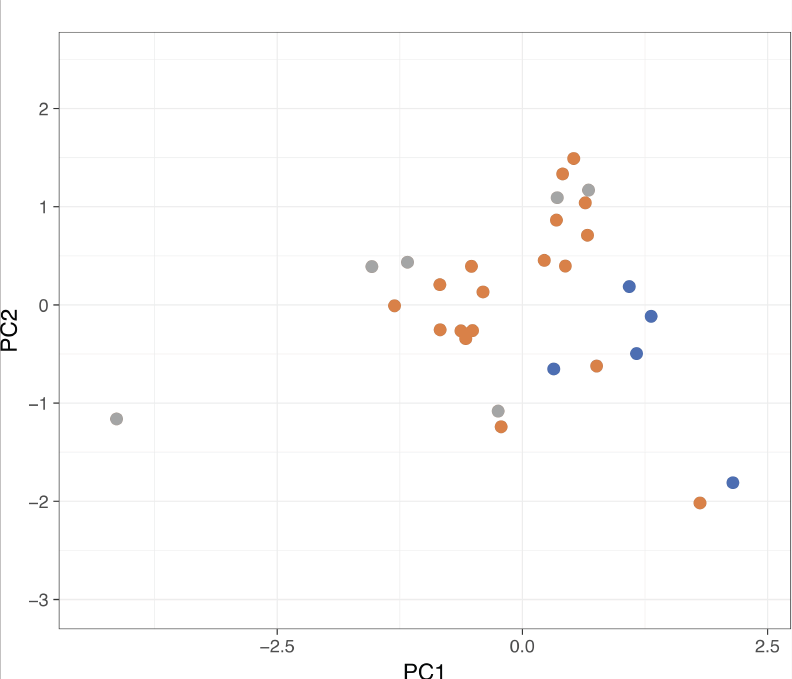
Principal component analysis of the CarrierSeq target reads (*n = 29*). There does not appear to be any separability based on read length, quality score, and DUST score parameters.

### 4.4. Detecting non-standard bases

Raw nanopore sequencing data represent the translocation of unique k-mers, or nucleotide sequences of k bases, through a critical pore region which produces an associated ionic current signal. For simplicity, let us imagine a sliding window that is 3 bases wide (a 3-mer) moving one base at a time over a single-stranded DNA of sequence AATCG. As the window slides we should observe AAT, ATC, and TCG each producing a unique signal. A moving 3-mer window over a DNA strand containing the four standard nucleobases (ATCG) could encounter up to 64 distinctive combination. Therefore, a DNA strand containing a theoretical fifth non-standard base would have 125 possible combinations and so on. Raw data is then basecalled by applying algorithms such as Hidden Markov Models (Simpson *et al.*, 2016) or Recurrent Neural Networks (Boža et al., 2017) that translate the collection of k-mers into consensus nucleic acid sequences. In previous work, we have demonstrated the capacity for these algorithms to detect non-standard bases by sequencing Poly(deoxyinosinic-deoxycytidylic) acid, a synthetic DNA polymer composed of alternating deoxy-inosine (I) and deoxy-cytosine (C) bases (Carr et al., 2017). This proof of principle demonstrates the detection of the non-standard inosine nucleoside, the nucleobase (hypoxanthine) of which which has been identified within meteorites (Martins et al., 2008); furthermore, work by others has demonstrated the ability to detect base modification such as methylation (Simpson et al., 2016) using novel basecalling algorithms.

## 5. Conclusions

Life on Mars, if it exists or existed in the not-so-distant past, may potentially be detected via in situ nucleic acid extraction and nanopore sequencing. We believe nucleic acids provide a sensitive and unambiguous indicator for life that facilitates distinguishing between forward contamination and putative nucleic acid-based Martian life (Isenbarger et al., 2008). Here, we have validated methodologies that enable the extraction and sequencing of nucleic acids from low-biomass Mars analog soils. Our results indicate that a ‘near universal’ extraction protocol may be capable of analyzing soils from various environments on Mars. Furthermore, nucleic acid—based life detection on Mars may only be practical in the context of extant or recently dead life-intact cells as extracellular DNA preserved in soil may be destroyed during sample preparation.

## Acknowledgements

This work was supported by NASA MATTISSE award NNX15AF85G and the MIT Shrock and OGE Diversity Graduate Fellowships.

## Abbreviations

SETG, Search for Extraterrestrial Genomes; DNA, deoxyribonucleic acid; gDNA, genomic deoxyribonucleic acids; ppb, parts per billion; RNA, ribonucleic acid; PCR, polymerase chain reaction; ddPCR, droplet digital polymerase chain reaction; NASA, National Aeronautics and Space Administration; HQNRs, high-quality noise reads.

